# Region-specific and state-dependent astrocyte Ca^2+^ dynamics during the sleep-wake cycle in mice

**DOI:** 10.1101/2020.11.16.385823

**Authors:** Tomomi Tsunematsu, Shuzo Sakata, Tomomi Sanagi, Kenji F. Tanaka, Ko Matsui

**Affiliations:** Super-network Brain Physiology, Graduate School of Life Sciences, Tohoku University, Sendai 980-8577, Japan; Advanced Interdisciplinary Research Division, Frontier Research Institute for Interdisciplinary Sciences, Tohoku University, Sendai 980-8578, Japan; Precursory Research for Embryonic Science and Technology, Japan Science and Technology Agency, Kawaguchi 332-0012, Japan; Strathclyde Institute of Pharmacy and Biomedical Sciences, University of Strathclyde, Glasgow G4 0RE, UK; Department of Neuropsychiatry, Keio University School of Medicine, Tokyo 160-8582, Japan

## Abstract

Neural activity is diverse, and varies depending on brain regions and sleep/wakefulness states. However, whether astrocyte activity differs between sleep/wakefulness states, and whether there are differences in astrocyte activity among brain regions remain poorly understood. In this study, we recorded astrocyte intracellular calcium (Ca^2+^) concentrations of mice during sleep/wakefulness states in the cortex, hippocampus, hypothalamus, cerebellum, and pons using fiber photometry. For this purpose, male transgenic mice in which their astrocytes specifically express the genetically encoded ratiometric Ca^2+^ sensor YCnano50 were used. We demonstrated that Ca^2+^ levels in astrocytes significantly decrease during Rapid Eye Movement (REM) sleep and increase after the onset of wakefulness. In contrast, differences in Ca^2+^ levels during non-Rapid Eye Movement (NREM) sleep were observed among different brain regions, and no significant decrease was observed in the hypothalamus and pons. Further analyses focusing on the transition between sleep/wakefulness states and correlation analysis with episode duration of REM showed that Ca^2+^ dynamics differed among brain regions, suggesting the existence of several clusters. To quantify region-specific Ca^2+^ dynamics, principal component analysis was performed to uncover three clusters; i.e., the first comprised the cortex and hippocampus, the second comprised the cerebellum, and the third comprised the hypothalamus and pons. Our study demonstrated that astrocyte Ca^2+^ levels change substantially according to sleep/wakefulness states. These changes were generally consistent, unlike neural activity. However, we also clarified that Ca^2+^ dynamics varies depending on the brain region, implying that astrocytes may play various physiological roles in sleep.

**Significance statement:** Sleep is an instinctive behavior of many organisms. In the previous five decades, the mechanism of the neural circuits controlling sleep/wakefulness states and the neural activities associated with sleep/wakefulness states in various brain regions have been elucidated. However, whether astrocytes, which are a type of glial cell, change their activity during different sleep/wakefulness states is poorly understood. Here, we demonstrated that dynamic changes in intracellular Ca^2+^ concentrations occur in the cortex, hippocampus, hypothalamus, cerebellum, and pons of genetically modified mice during natural sleep. Further analyses demonstrated that Ca^2+^ dynamics slightly differ among different brain regions, implying that the physiological roles of astrocytes in sleep/wakefulness might vary depending on the brain region.

## Introduction

Astrocytes, which are the principal subtype of glial cells, are essential for central nervous system development and function. Many previous studies have clarified that astrocytes have housekeeping roles in brain function, contributing to ion and neurotransmitter homeostasis, formation and maintenance of the blood-brain barrier (Sofroniew and Vinters, 2010; Bojarskaite et al., 2020), regulation of blood flow, trophic, antioxidant and metabolic support for neurons (Magistretti and Allaman, 2018), neurotransmitter recycling (Sofroniew and Vinters, 2010), and regulation of synaptogenesis and synaptic transmission (Allen, 2014; Allen and Eroglu, 2017). All these physiological functions of astrocytes are strongly associated with the dynamics of their intracellular calcium (Ca^2+^) concentration. Intrinsic signals, including those involving neurotransmitters, protons, cannabinoids, polyphosphate, and endothelin result in increases in astrocyte Ca^2+^ concentrations (Gourine et al., 2010; Navarrete and Araque, 2010; Filosa et al., 2012; Min and Nevian, 2012; Holmstrom et al., 2013). In turn, the activated astrocytes release neuroactive substances called gliotransmitters, which activate neurons and vascular smooth muscle (Sasaki et al., 2012; Araque et al., 2014; Beppu et al., 2014; Savtchouk and Volterra, 2018).

In addition, it has become clear that astrocytes are extensively involved in the regulation and physiological functions of mammalian sleep (Frank, 2019). There is evidence that astrocytes regulate sleep pressure through soluble NSF attachment protein receptor (SNARE)-dependent adenosine release (Halassa et al., 2009; Florian et al., 2011). Astrocytes also promote the sleep-dependent clearance of brain waste products, such as β-amyloid (Xie et al., 2013). The dysfunction of gap junctions in astrocytes leads to the inability to transfer lactate to wake-promoting orexin neurons in the lateral hypothalamic area, resulting in a sleep disorder (Clasadonte et al., 2017), whereas optogenetic stimulation of astrocytes in the posterior hypothalamus increases sleep (Pelluru et al., 2016). In addition, astrocytes can also regulate cortical state switching (Poskanzer and Yuste, 2016). These results strongly support that astrocytes play a pivotal role in sleep as well as other brain functions.

However, astrocytic Ca^2+^ dynamics in natural sleep has been poorly understood, although an imaging study demonstrated that general anesthesia disrupts astrocyte Ca^2+^ signaling in mice (Thrane et al., 2012). Recent studies have recorded astrocyte Ca^2+^ changes during sleep/wakefulness in mice, but to date, data has been limited to that of only the cortex (Bojarskaite et al., 2020; Ingiosi et al., 2020). Accumulating evidence shows that astrocytes are heterogeneous with respect to their transcriptomes and functions among various brain regions as well as among various types of neurons (Chai et al., 2017; Morel et al., 2017; Zeisel et al., 2018; Batiuk et al., 2020; Bayraktar et al., 2020; Lozzi et al., 2020). It is well known that neurons in various brain regions control the sleep/wakefulness state, as well as rapid eye movement (REM)/ non-REM (NREM) sleep, via a flip-flop circuit. It has been reported that the hypothalamus, midbrain, and brainstem play important roles in the brain (Sakurai, 2007; Weber and Dan, 2016; Scammell et al., 2017; Liu and Dan, 2019). However, little is known about how astrocyte Ca^2+^ concentrations in brain regions other than the cortex change depending on the sleep/wakefulness state, and whether there are dynamic differences in astrocyte Ca^2+^ levels among brain regions. Therefore, in this study, we performed fiber photometry recordings of astrocyte Ca^2+^ dynamics from the cerebellum, cortex, hippocampus, hypothalamus, and pons of mice during the sleep-wake cycle. To optically record astrocyte Ca^2+^ dynamics, we used *megalencephalic leukoencephalopathy with subcortical cysts 1* (*Mlc1*)-tTA; TetO-YCnano50 bigenic mice (Horikawa et al., 2010; Tanaka et al., 2010; Tanaka et al., 2012; Kanemaru et al., 2014).

In this study, we demonstrated that Ca^2+^ levels in astrocytes consistently decrease during REM sleep, and immediately increase with the onset of wakefulness. In contrast, differences in Ca^2+^ levels during NREM sleep were observed in various brain regions. There was a significant decrease in Ca^2+^ levels in the cerebellum, cortex, and hippocampus during NREM sleep, but no significant decrease was observed in the hypothalamus and pons. Further detailed analyses also demonstrated that there are substantial differences in Ca^2+^ dynamics among the various brain regions.

## Materials and Methods

### Animals

All experimental procedures involving animals were approved by the Animal Care and Use Committee of Tohoku University (approval no.: 2019LsA-018) and were conducted in accordance with the National Institute of Health guidelines. All efforts were made to minimize animal suffering and discomfort, and to reduce the number of animals used. *Mlc1*-tTA; TetO-YCnano50 mice, which were used to monitor the dynamics of intracellular calcium concentration in astrocytes, were produced by crossing *Mlc1-tTA* mice (Tanaka et al., 2010) with TetO-YCnano50 mice (Kanemaru et al., 2014). The following PCR primer sets were used for mouse genotyping: MlcU-675 (5’-AAATTCAGGAAGCTGTGTGCCTGC-3’) and mtTA24L (5’-CGGAGTTGATCACCTTGGACTTGT-3’) for *Mlc1*-tTA mice; and tetO-up (5’-AGCAGAGCTCGTTTAGTGAACCGT-3’) and intronlow (5’-AAGGCAGGATGATGACCAGGATGT-3’) for TetO-YCnano50 mice. Mice were housed under a controlled 12 h/12 h light/dark cycle (light on hours: 8:30–20:30). Mice had *ad libitum* access to food and water. A total of ten male mice (seven *Mlc1*-tTA; TetO-YCnano50 mice and three *Mlc1*-tTA mice as controls) were used in this study. The following number of mice were used for the recording of each brain area: cerebellum, three mice (5 recordings); cortex, three mice (6 recordings); hippocampus, one mouse (2 recordings); hypothalamus, two mice (4 recordings); pons, one mouse (2 recordings). For the recording of control *Mlc1*-tTA mice, three mice were used (8 recordings).

### Surgical procedures

Male *Mlc1*-tTA;TetO-YCnano50 bigenic mice (≥ 12 weeks of age) were used. Stereotaxic surgery was performed under anesthesia with pentobarbital (5 mg/kg, intraperitoneal (i.p.) injection as induction) and with isoflurane (1%–2% for maintenance) using a vaporizer for small animals (Bio Research Center) with the mice positioned in a stereotaxic frame (Narishige). Two bone screws were implanted on the skull as electrodes for cortical electroencephalograms (EEGs) and twisted wires (AS633, Cooner Wire) were inserted into the neck muscle as an electrode for electromyogram (EMG). Another bone screw was implanted in the cerebellum as a ground. All electrodes were connected to a pin socket.

For fiber photometry experiments, a cannula (CF440-10, Thorlabs) with a glass optical fiber (φ 400 μm, 0.39 NA, Thorlabs) was implanted into the cortex (1.2 mm posterior, 3.1 mm lateral from bregma, 0.3 mm depth from the brain surface), hippocampus (1.7 mm posterior, 1.5 mm lateral from bregma, 1.3 mm depth from the brain surface), hypothalamus (1.8 mm posterior, 1.0 mm lateral from bregma, 4.5 mm depth from the brain surface), cerebellum (6.0 mm posterior, 1.0 mm lateral from bregma, and 0.5 mm depth from the brain surface), and pons (5.1 mm posterior, 1.2 mm lateral from bregma, and 3.5 mm depth from the brain surface). All electrodes and optical fiber cannula were fixed to the skull with dental cement. To fixate the head of mice, a stainless chamber frame (CF-10, Narishige) was also attached to the skull using dental cement. After the surgery, the mice were left to recover for at least 5 days. During the habituation period, mice were placed in a head-fixed apparatus (MAG-1, Narishige), by securing them by the stainless chamber frame and placing them into an acrylic tube. This procedure was continued for at least 5 days, during which the duration of head-fixation was gradually extended from 10 to 120 min.

### *In vivo* fiber photometry experiments in the head-fixed condition

To detect the dynamics of intracellular Ca^2+^ concentrations in astrocytes, a fiber photometric system (Lucir) was used. A 420-nm violet light emitting diode (LED) (Doric) was used to obtain Ca^2+^-dependent signals. Recording in the head-fixed condition was performed for about five hrs a day. During recording, EEG and EMG were recorded continuously, whereas excitation light (20 Hz, 5 msec in width) was intermittently illuminated at random for 4 minutes each time. The input light was reflected off a dichroic mirror (FF458-Di02, Semrock) coupled to an optical fiber. LED power was 1.12 ± 0.18 mW/mm^2^ at the fiber tip. Light emission of cyan and yellow fluorescence from YCnano50 was collected via an optical fiber cannula, divided by a dichroic mirror (FF509-FDi01, Semrock) into cyan (483/32 nm bandpass filter, Semrock) and yellow (542/27 nm bandpass filter, Semrock), and detected by each photomultiplier (Lucir). Excitation signals were generated by a pulse generator (AWG-50, Elmos) to control the LEDs. Fluorescence data were acquired at a sampling rate of 1 kHz through an analog-to-digital converter (Micro1401-3, Cambridge Electronic Design (CED)). At the same time as the fluorescence recording, EEG and EMG signals were amplified (DAM50, World Precision Instruments), filtered, and digitized at 1 kHz using an analog-to-digital converter. EEG and EMG signals were high-pass- and low-pass-filtered at 0.1 Hz and 300 Hz, respectively. Data were recorded using Spike2 software (CED).

### Histological analysis

To confirm the position of the implanted fiber optics, mice were deeply anesthetized with isoflurane and perfused sequentially with 20 mL of chilled saline and 20 mL of chilled 4% paraformaldehyde in phosphate buffer solution (Nacalai Tesque). The brains were removed and immersed in the above fixation solution overnight at 4 °C, and then immersed in 30% sucrose in phosphate-buffered saline (PBS) for at least 2 days. The brains were quickly frozen in embedding solution (Sakura Finetek) and cut into coronal sections using a cryostat (CM3050, Leica) at a thickness of 50 μm. The brain sections were mounted on APS-coated slides and coverslipped with 50% glycerol in PBS. The sections were observed using a fluorescence microscope (BZ-9000, Keyence).

### Data analysis

#### Sleep scoring

Polysomnographic recordings were automatically scored offline, with each epoch scored as wakefulness, NREM sleep, or REM sleep by SleepSign (KISSEI COMTEC), in 4 sec epochs, according to standard criteria (Radulovacki et al., 1984; Tobler et al., 1997). All vigilance state classifications assigned by SleepSign were confirmed visually. The same individual, blinded to mouse genotype and experimental condition, scored all EEG/EMG recordings. Spectral analysis of the EEG was performed by fast Fourier transform, which yielded a power spectral profile with a 1 Hz resolution divided into delta (1-5 Hz), theta (6-10 Hz), alpha (10-13 Hz), beta (13-25 Hz), and gamma (30-50 Hz) waves. To quantify EMG amplitude, the root-mean-square (rms) was calculated.

#### Fiber photometry signal processing

In Figures 1, 2, and 3, axoGraph was used to calculate yellow fluorescence protein (YFP) to cyan fluorescence protein (CFP) (Y/C) ratios. The average value of the YFP intensity and CFP intensity for each light illumination (5 msec) was calculated, and then the Y/C ratio was calculated. For the comparison of Y/C ratios during the sleep/wakefulness states, the Y/C ratio of each episode was normalized with the average value during wakefulness set as 1. For sleep/wakefulness state transition analyses, four consecutive epochs (16 sec) of one state followed immediately by eight consecutive epochs (32 sec) of a distinct state were used. To assess correlations between Y/C ratio and EEG/EMG power (Figure 2), spectral densities of EEG signals in every 1-sec window were estimated using a multitaper method (http://chronux.org/).

**Figure 1.**
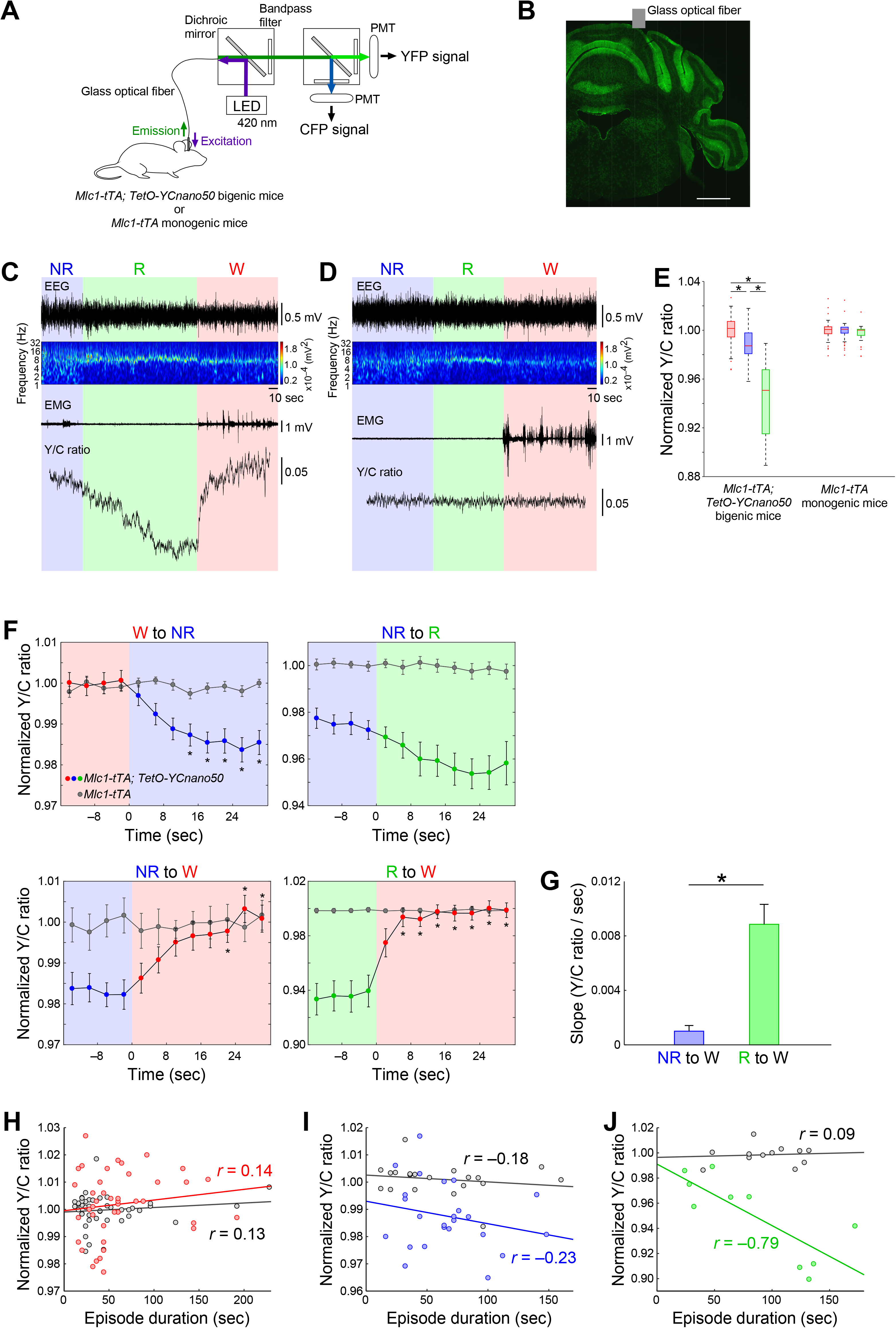
Astrocyte Ca^2+^ dynamics in the cerebellum during sleep/wakefulness states. (A) Schematic drawing showing the fiber photometry system used in this study. Fluorescence emission is applied from the LED light source. Yellow and cyan fluorescence signals are corrected by bandpass filters and enhanced by photomultipliers (PMT). (B) The location of the glass optical fiber, which is implanted in the cerebellum of *Mlc1*-tTA; TetO-YCnano50 bigenic mice (6.0 mm posterior, 1.0 mm lateral from bregma, 0.5 mm depth from the brain surface). Scale bar = 1 mm. (C and D) Representative traces of EEG, EEG power density spectrum, EMG, and cerebellar astrocyte Ca^2+^ signals (Y/C ratio) in *Mlc1*-tTA; TetO-YCnano50 bigenic mice (C) and *Mlc1*-tTA monogenic mice (D). (E) Box plots summarizing the data from C and D. Y/C ratios were normalized to the value of each episode, with the average value of awakening set as 1. \**, p* < 0.05. (F) Y/C ratios during the transitions between the sleep/wakefulness states. Transitions occurred at time 0. Data are from 4-sec intervals characterized by state transitions. The line graph with the colored circles and gray circles are a summary of the data from *Mlc1*-tTA; TetO-YCnano50 bigenic mice and *Mlc1*-tTA monogenic mice, respectively. \**, p* < 0.05 vs. the fourth epoch immediately before state transition. (G) The bar graph represents the slope of the Y/C ratio when awakening from NREM sleep and REM sleep. \**, p* < 0.05. (H, I, and J) Analyses of the correlation between Y/C ratio and episode duration of wakefulness (H), NREM sleep (I), and REM sleep (J). Colored circles and gray circles indicate the summary of data from *Mlc1*-tTA; TetO-YCnano50 bigenic mice and *Mlc1*-tTA monogenic mice, respectively. NR, NREM sleep; R, REM sleep; W, wakefulness. Values are shown as means ± SEM.

**Figure 2.**
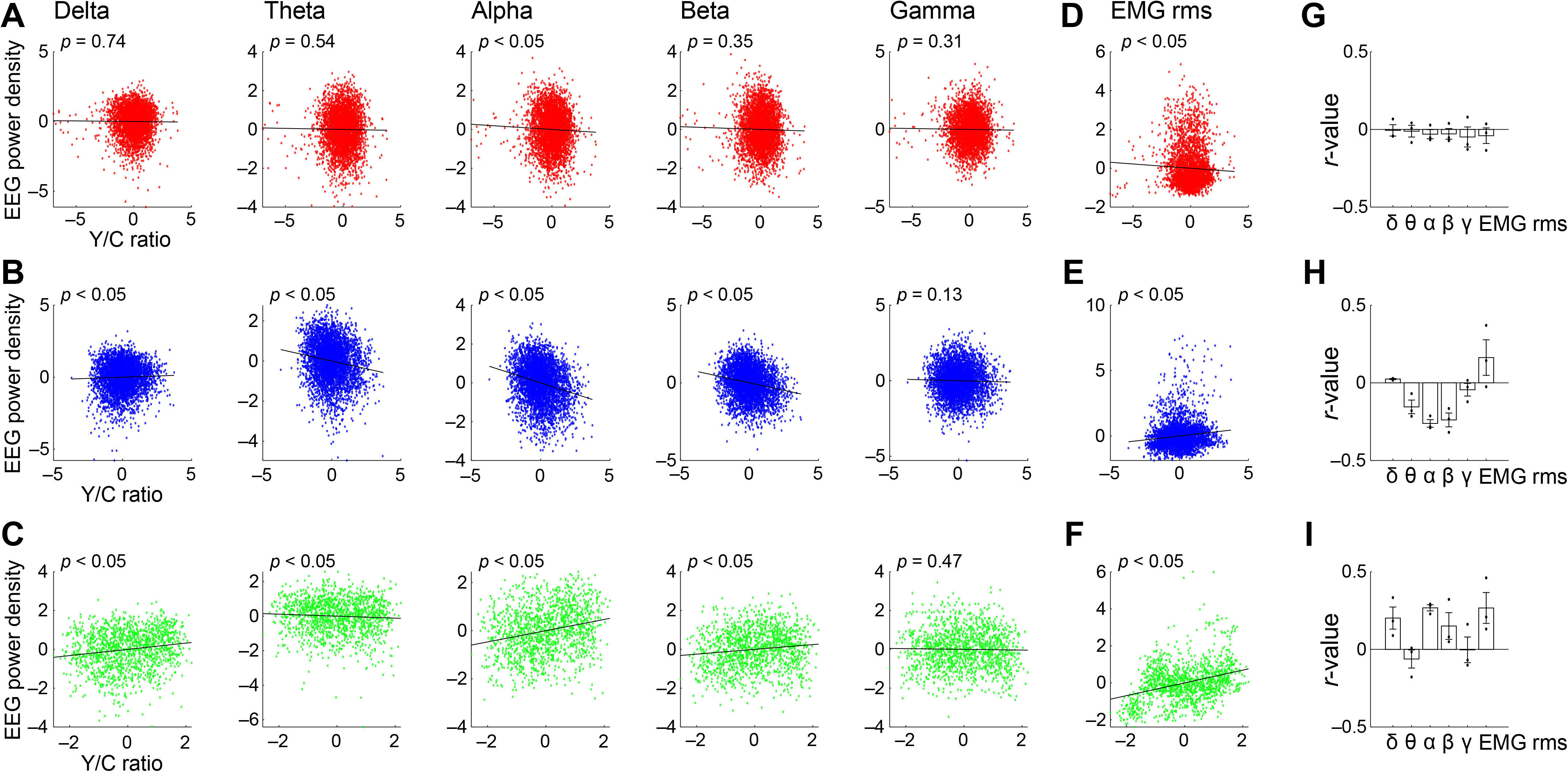
Correlation between EEG/EMG and cerebellar astrocytic Ca^2+^ signals during different sleep/wakefulness states. (A, B, and C) Correlation analyses between normalized (z-scored) cerebellar astrocytic Y/C ratios and normalized (z-scored) EEG power densities in the delta (1–5 Hz), theta (6–10 Hz), alpha (10–13 Hz), beta (13–25 Hz), and gamma (30–50 Hz) wave during wakefulness (A), NREM sleep (B), and REM sleep (C). (D, E, and F) Correlation analyses between normalized Y/C ratios and normalized root-mean-squares (rms) of EMG during wakefulness (D), NREM sleep (E), and REM sleep (F). The data in this figure were analyzed with a 1 sec bin size. (G, H, and I) Bar graphs showing correlation coefficients summarizing the data from A to F. The correlation coefficient of each recording was cross-validated by splitting the data into first and second halves.

**Figure 3.**
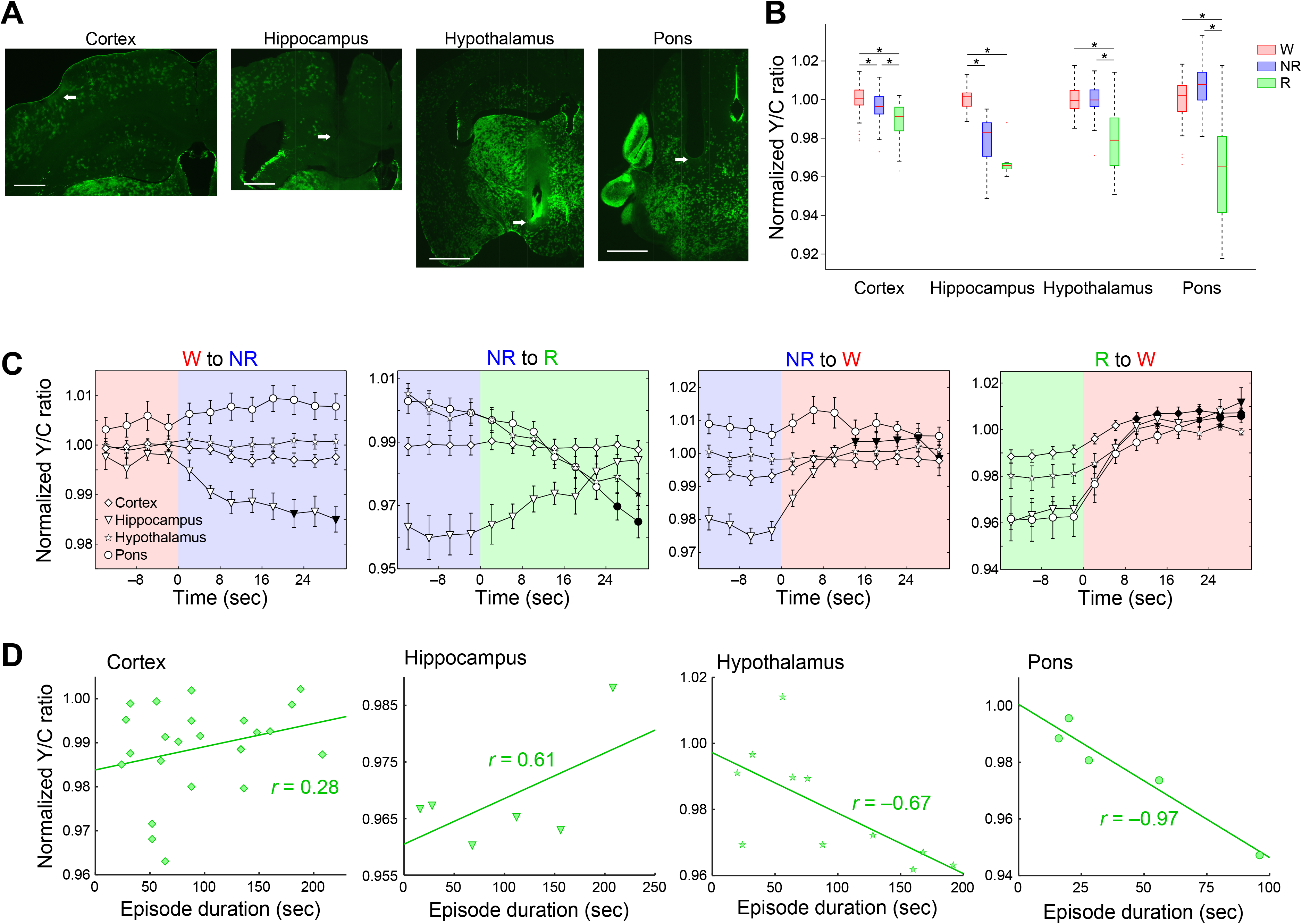
Astrocyte Ca^2+^ dynamics during different sleep/wakefulness states in various brain regions. (A) The location of the glass optical fiber, which is implanted in the cortex, hippocampus, hypothalamus, and pons, respectively. Arrow indicate the tip of the optical fiber. Cortex, hippocampus; Scale bar = 500 μm. Hypothalamus, pons; Scale bar = 1 mm. (B) Box plots summarizing the data of normalized Y/C ratios obtained from the cortex, hippocampus, hypothalamus, and pons. *, *p* < 0.05. (C) Y/C ratios for the transition of sleep/wakefulness states in multiple brain regions. Black filled symbols indicate *p* < 0.05 vs. the fourth epoch immediately before state transition in each brain region. (D) Correlation analyses between episode durations of REM sleep and Y/C ratios in the cortex, hippocampus, hypothalamus, and pons. NR, NREM sleep; R, REM sleep; W, wakefulness. Values are represented as means ± SEM.

In Figure 4, all data analyses were performed by custom written MATLAB software (MathWorks). To compute normalized Y/C ratios during the time-normalized episodes, Y/C ratios were first normalized by the mean Y/C ratio during wakefulness. To obtain the time-normalized Ca^2+^ dynamics, each episode was segmented into 5 bins and the mean Y/C ratio was computed for each bin. Episodes were classified based on vigilance states before and after the episode.

**Figure 4.**
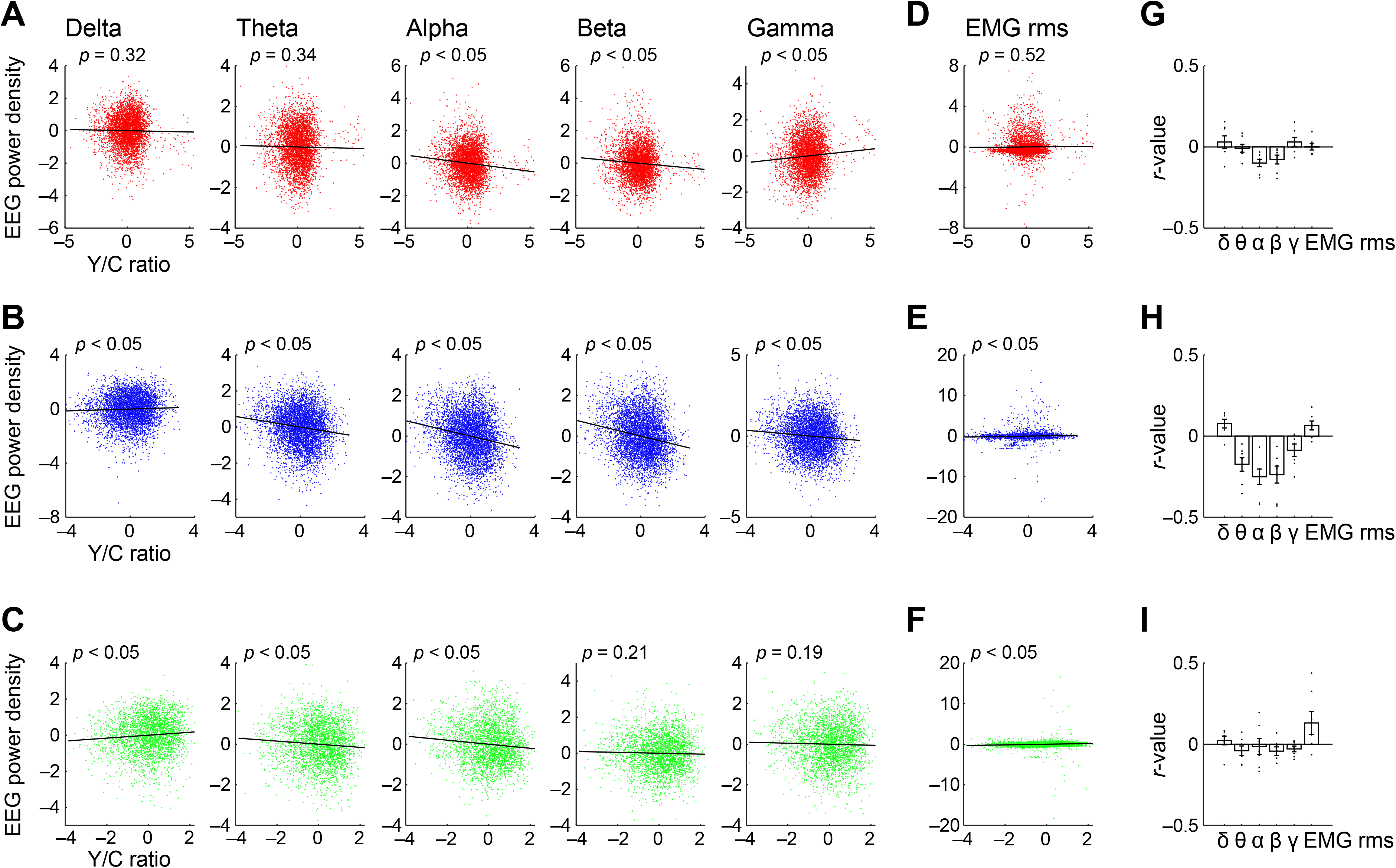
Correlation of EEG/EMG and cortical astrocyte Ca^2+^ signals during different sleep/wakefulness states. (A, B, and C) Correlation analyses between normalized (z-scored) Y/C ratios from the cortex and normalized (z-scored) EEG power densities in the delta (1–5 Hz), theta (6–10 Hz), alpha (10–13 Hz), beta (13–25 Hz), and gamma (30–50 Hz) wave during wakefulness (A), NREM sleep (B), and REM sleep (C). (D, E, and F) Correlation analyses between normalized Y/C ratios and normalized root-mean-squares (rms) of EMG during wakefulness (D), NREM sleep (E), and REM sleep (F). The data in this figure were analyzed with a 1 sec bin size. (G, H, and I) Bar graphs showing correlation coefficients summarizing the data from A to F. The correlation coefficient of each recording was cross-validated by splitting the data into first and second halves.

#### Principal component analysis (PCA) and decoding

To perform PCA, Ca^2+^ dynamics of each mouse was represented as a single vector. Each vector was composed of the mean profile of the normalized Y/C ratio throughout state transitions (6 bins/transition type) and time-normalized episodes (5 bins/state history). The state transitions included the transitions from Wakefulness to NREM, from NREM to REM, and from REM to wakefulness. Episodes that lasted for at least 12 sec (three epochs) were included. As a result, the data matrix was composed of recordings from three cortices, one hippocampus, two hypothalamus, one pons, two cerebella, and four controls.

To decode sleep/wakefulness states based on Ca^2+^ signals, the same approach as described previously was used (Tsunematsu et al., 2020). Briefly, the mean of Y/C ratio was computed in each corresponding window (4 sec). After training a linear classifier, classification performance was calculated with 4-fold cross validation.

#### Experimental design and statistical analysis

Data are presented as the mean ± SEM unless otherwise stated. Statistical analyses were performed using MATLAB. Multiple group comparisons were performed by one-way analysis of variance (ANOVA) in samples with a Gaussian distribution and by the Kruskal-Wallis test in samples with a non-Gaussian distribution, with the *post-hoc* Bonferroni test. In Figure 4D, one-way ANOVA with the *post-hoc* Tukey’s honest significance difference (HSD) test was performed. *P*-values of less than 0.05 were considered to indicate a statistically significant difference between groups.

## Results

### State-dependent Ca^2+^ dynamics in cerebellar astrocytes

To elucidate the dynamics of intracellular Ca^2+^ concentration in astrocytes during sleep/wakefulness states, we used *Mlc1*-tTA; TetO-YCnano50 bigenic mice. Mlc1 is an astrocyte-specific protein with unknown function, which is highly expressed in perivascular astrocyte end-feet and astrocyte-astrocyte contacts (Boor et al., 2005; Teijido et al., 2007). Thus, *Mlc1*-tTA; TetO-YCnano50 bigenic mice specifically express the ratiometric Ca^2+^ indicator YCnano50 in astrocytes (Kanemaru et al., 2014). In this study, fiber photometry was used to record changes in astrocyte Ca^2+^ concentrations from head-fixed mice, as Y/C ratios (Fig. 1A). To determine the sleep/wakefulness state, EEG and EMG electrodes were implanted. We first focused on cerebellar astrocytes and investigated the changes in Y/C ratios. It has been reported that the activity of Bergmann glial cells, a specific type of radial astrocyte in the cerebellum, is inhibited in the anesthetized state compared with in the awake state (Hoogland et al., 2009; Nimmerjahn et al., 2009). Therefore, we hypothesized that cerebellar astrocytes may demonstrate state-dependent Ca^2+^ dynamics throughout the sleep-wake cycle. To this end, we took advantage of the strong fluorescence intensity of cerebellar astrocytes in this bigenic mouse line (Kanemaru et al., 2014).

To deliver excitation light and collect fluorescence signals from the cerebellum, a glass optical fiber was implanted in the cerebellum of mice (Fig. 1B). Ca^2+^ signal dynamics were observed during the light period (9:00 to 15:00). Excitation light (20 Hz, 5 msec in width) was intermittently illuminated at random for 4 minutes. Y/C ratios gradually decreased during sleep, showing the lowest value during REM sleep, and instantaneously increased with awakening in *Mlc1*-tTA; TetO-YCnano50 mice (Fig. 1C). In contrast, Y/C ratios did not change with sleep/wakefulness state in control *Mlc1*-tTA mice (Fig. 1D), indicating that no intrinsic signals were detected in this experimental condition. Normalized Y/C ratios in wakefulness, NREM sleep, and REM sleep in cerebellar astrocytes of *Mlc1*-tTA; TetO-YCnano50 mice were 1.000 ± 0.001 (n = 66 episodes from 5 recording sessions and 3 animals), 0.988 ± 0.002 (n = 63 episodes from 5 recording sessions and 3 animals), and 0.944 ± 0.008 (n = 17 episodes from 3 recording sessions and 2 animals), respectively (Fig. 1E) (Kruskal-Wallis: F(2, 143) = 56.68, *p* < 0.05; followed by multiple comparison with the Bonferroni test: *, *p* < 0.05). In contrast, normalized Y/C ratios during wakefulness, NREM sleep, and REM sleep in cerebellar astrocytes of *Mlc1*-tTA mice were 1.000 ± 0.001 (n = 60 episodes from 5 recording sessions and 3 animals), 1.000 ± 0.001 (n = 70 episodes from 5 recording sessions and 3 animals), and 0.997 ± 0.001 (n = 33 episodes from 5 recording sessions and 3 animals), respectively (Fig. 1E) (Kruskal-Wallis: F(2, 160) = 4.04, *p* = 0.13, no significant difference [NS]). These results indicate that Ca^2+^ concentrations of cerebellar astrocytes change substantially with sleep/wakefulness state in mice.

We next focused on Ca^2+^ dynamics during the transition between sleep/wakefulness states (Fig. 1F). Y/C ratios gradually decreased after the onset of NREM sleep following wakefulness. Sixteen seconds after the start of NREM sleep, Y/C ratios significantly decreased compared with when mice were awake (n = 26 episodes from 3 recording sessions and 2 animals) (one-way ANOVA: F(11, 300) = 6.79, *p* < 0.05; followed by multiple comparisons with the Bonferroni test: *, *p* < 0.05 vs the fourth epoch immediately before state transition). A slow decrease in Y/C ratios was also observed in the transition from NREM sleep to REM sleep, but there was no significant difference (n = 10 episodes from 2 recording sessions and 2 animals) (one-way ANOVA: F(11, 108) = 2.35, *p* < 0.05; followed by multiple comparison with the Bonferroni test: *p* ≥ 0.05, NS). During the transition from NREM and REM sleep to wakefulness, the Y/C ratio increased (From NREM to wake; n = 19 episodes from 4 recording sessions and 3 animals. From REM to wake; n = 14 episodes from 3 recording sessions and 2 animals) (From NREM to wake, one-way ANOVA: F(11, 216) = 5.47, *p* < 0.05; followed by multiple comparisons with the Bonferroni test: *, *p* < 0.05 vs the fourth epoch immediately before state transition. From REM to wake, one-way ANOVA: F(11, 156) = 12.87, *p* < 0.05; followed by multiple comparisons with the Bonferroni test: *, *p* < 0.05 vs the fourth epoch immediately before state transition). However, the slopes were completely different between from NREM to wakefulness and from REM to wakefulness. The slope was calculated by dividing the difference in Y/C ratios immediately before and after the transition by four seconds. The slope from REM to wakefulness (n = 14 episodes from 3 recording sessions and 2 animals) was significantly larger than the slope from NREM to wakefulness (Fig. 1G) (n = 19 episodes from 4 recording sessions and 3 animals) (unpaired *t*-test: *, *p* < 0.05). These results indicate that the signaling pathway that increases intracellular Ca^2+^ concentrations might differ between from REM sleep and from NREM sleep, although they do not appear to precede the four-second bin.

Next, correlation analysis was performed to investigate the association between episode duration of wakefulness, NREM, and REM, and changes in Ca^2+^ concentrations (Fig. 1H and 1J). There was no significant correlation between wakefulness and NREM sleep in either *Mlc1*-tTA; TetO-YCnano50 mice (Wakefulness; n = 41 episodes from 3 recording sessions and 2 animals. NREM; n = 24 episodes from 3 recording sessions and 2 animals) (wakefulness; *r* = 0.14, *p* = 0.38, NS. NREM; *r* = −0.23, *p* = 0.28, NS) nor in *Mlc1*-tTA mice (Wakefulness; n = 46 episodes from 5 recording sessions and 3 animals. NREM; n = 20 episodes from 4 recording sessions and 3 animals) (wakefulness; *r* = 0.13, *p* = 0.40, NS. NREM; *r* = −0.18, *p* = 0.45, NS). In contrast, there was a significant negative correlation between the episode duration of REM and Y/C ratio in *Mlc1*-tTA; TetO-YCnano50 mice (Fig. 1J) (n = 10 episodes from 2 recording sessions and 2 animals) (*r* = −0.79, *p* < 0.05), which is in good agreement with the gradual decrease in Ca^2+^ level during REM sleep. On the other hand, no significant correlation was observed in *Mlc1*-tTA mice (Fig. 1J) (n = 12 episodes from 2 recording sessions and 2 animals) (*r* = 0.09, *p* = 0.79, NS). These results indicate that Ca^2+^ concentration in cerebellar astrocytes decreases as REM sleep episode duration increases.

### Correlations between EEG/EMG and Ca^2+^ signals in cerebellar astrocytes during sleep/wakefulness

We demonstrated that cerebellar astrocyte Ca^2+^ concentrations change dynamically with sleep/wakefulness state. Therefore, we next investigated whether Ca^2+^ fluctuations in cerebellar astrocytes also correlate with electrophysiological features during each sleep/wakefulness state. For this purpose, we investigated the association between Y/C ratios and cortical EEGs and EMGs during wakefulness, NREM, and REM sleep (50 recording sessions and 3 animals). Cortical EEGs were analyzed by dividing them into delta (1–5 Hz), theta (6–10 Hz), alpha (10–13 Hz), beta (13–25 Hz), and gamma (30–50 Hz) wave components. Regarding EMGs, their magnitudes were evaluated by calculating the rms. Then, their correlation with Y/C ratios was analyzed. During wakefulness, although Y/C ratios showed a significant negative correlation with alpha waves and EMG rms (Fig. 2A and 2D), the effect was weak (Fig. 2G), suggesting little cofluctuation of Ca^2+^ and electrophysiological signals during wakefulness. During NREM sleep, Y/C ratios were positively correlated with delta waves and EMG rms, and negatively correlated with theta, alpha, and beta waves (Fig. 2B, 2E, and 2H). During REM sleep, however, Y/C ratios demonstrated a positive correlation with delta, alpha, and beta waves and EMG rms, and a negative correlation with theta waves (Fig. 2C, 2F, and 2I). Although this correlation analysis demonstrated significant differences, no highly positive nor highly negative correlation was identified. A minor but statistically significant correlation between cerebellar astrocyte Ca^2+^ concentrations and electrophysiological features, particularly during sleep, was identified.

### State-dependent astrocyte Ca^2+^ dynamics among various brain regions

We next assessed whether state-dependent changes in astrocyte Ca^2+^ concentrations are observed not only in the cerebellum but also in other brain regions, and whether the dynamics differ depending on the brain region. Fiber photometry recordings were performed using a glass optical fiber from the cortex and hippocampus, which have been reported to show diversity in neural activity corresponding to the sleep/wakefulness state (Vyazovskiy et al., 2009; Grosmark et al., 2012; Watson et al., 2016; Niethard et al., 2017), and from the hypothalamus and pons, which play a crucial role in the regulation of sleep/wakefulness (Sakurai, 2007; Tsunematsu et al., 2014; Hayashi et al., 2015; Weber et al., 2015; Weber and Dan, 2016; Scammell et al., 2017), using *Mlc1*-tTA; TetO-YCnano50 mice. To deliver excitation light and collect fluorescence signals, a glass optical fiber was implanted (Fig. 3A). Interestingly, intracellular Ca^2+^ levels in astrocytes dynamically changed depending on the sleep/wakefulness state throughout the brain regions that were monitored, and significantly decreased in all areas during REM sleep (Fig. 3B) (n = 69 (W), 68 (NR) and 25 (R) episodes from 5 recording sessions and 3 animals in the cortex. n = 23 (W), 24 (NR) and 8 (R) episodes from 2 recording sessions and 1 animal in the hippocampus. n = 60 (W), 69 (NR) and 20 (R) episodes from 4 recording sessions and 2 animals in the hypothalamus. n = 40 (W), 43 (NR) and 34 (R) episodes from 2 recording sessions and 1 animal in the pons) (Kruskal-Wallis: F(2, 159) = 29.3, *p* < 0.05 in the cortex. Kruskal-Wallis: F(2, 52) = 39.1, *p* < 0.05 in the hippocampus. Kruskal-Wallis: F(2, 146) = 30.0, *p* < 0.05 in the hypothalamus. Kruskal-Wallis: F(2, 114) = 55.8, *p* < 0.05 in the pons; followed by multiple comparisons by the Bonferroni test: *, *p* < 0.05.). However, significant decreases in Y/C ratios during NREM sleep were observed in the cortex and hippocampus compared with that during wakefulness. In contrast, in the hypothalamus and pons, no significant differences were observed between the Y/C ratios during wakefulness and NREM sleep, but Y/C ratios were significantly reduced during REM sleep compared with during NREM sleep. These results suggest that astrocyte Ca^2+^ levels are at a minimum during REM sleep whereas they are high during wakefulness. This trend is consistent with that observed in the cerebellum. On the other hand, Ca^2+^ dynamics during NREM sleep vary depending on the brain region.

Further analyses during the sleep/wakefulness state transition clarified the variety of Ca^2+^ dynamics among different brain regions (Fig. 3C). Focusing on the transition from wakefulness to NREM sleep, no significant changes were seen in the cortex, hypothalamus, and pons (n = 44 episodes from 6 recording sessions and 3 animals in the cortex. n = 38 episodes from 2 recording sessions and 1 animal in the hypothalamus. n = 17 episodes from 1 recording sessions and 1 animal in the pons) (Kruskal-Wallis: F(11, 516) = 11.7, *p* = 0.39, NS in the cortex. Kruskal-Wallis: F(11, 444) = 3.2, *p* = 0.99, NS in the hypothalamus; Kruskal-Wallis: F(11, 192) = 7.3, *p* = 0.77, NS in the pons), but Y/C ratios gradually decreased in the hippocampus and significantly after 24 seconds from state transition (n = 12 episodes from 2 recording sessions and 1 animal in the hippocampus) (Kruskal-Wallis: F(11, 132) = 49.2, p < 0.05; followed by multiple comparisons with the Bonferroni test: *p* < 0.05 vs the fourth epoch immediately before state transition in the hippocampus). A slow decrease Y/C ratios in the hypothalamus and pons was observed from NREM to REM sleep (n = 6 episodes from 2 recording sessions and 1 animal in the hypothalamus. n = 8 episodes from 2 recording sessions and 1 animal in the pons) (Kruskal-Wallis: F(11, 60) = 39.9, *p* < 0.05 in the hypothalamus; Kruskal-Wallis: F(11, 84) = 57.8, *p* < 0.05 in the pons, followed by multiple comparisons with the Bonferroni test: *p* < 0.05 vs the fourth epoch immediately before state transition), whereas it was almost constant in the cortex but was increased in the hippocampus (n = 15 episodes from 6 recording sessions and 3 animals in the cortex. n = 5 episodes from 2 recording session and 1 animal in the hippocampus) (Kruskal-Wallis: F(11, 168) = 0.98, *p* = 0.99, NS in the cortex; Kruskal-Wallis: F(11, 48) = 23.4, *p* < 0.05, followed by multiple comparisons with the Bonferroni test: NS vs the fourth epoch immediately before state transition in the hippocampus). At the time of transition from NREM sleep to wakefulness, Y/C ratios increased in the hippocampus (n = 12 episodes from 2 recording sessions and 1 animal in the hippocampus) (Kruskal-Wallis: F(11, 132) = 89.3, *p* < 0.05, followed by multiple comparisons with the Bonferroni test: *p* < 0.05 vs the fourth epoch immediately before state transition in the hippocampus), but there was no significant difference in the cortex, hypothalamus, and pons (n = 34 episodes from 7 recording sessions and 3 animals in the cortex. n = 25 episodes from 2 recording sessions and 1 animal in the hypothalamus. n = 7 episodes from 1 recording session and 1 animal in the pons) (Kruskal-Wallis: F(11, 396) = 31.8, *p* < 0.05, followed by multiple comparisons with the Bonferroni test: NS vs the fourth epoch immediately before state transition in the cortex. Kruskal-Wallis: F(11, 288) = 6.0, *p* = 0.87, NS in the hypothalamus. Kruskal-Wallis: F(11, 72) = 5.4, *p* = 0.91, NS in the pons). There were consistent increases immediately after the state change from REM sleep to wakefulness in all brain regions that were monitored (n = 20 episodes from 6 recording sessions and 3 animals in the cortex. n = 6 episodes from 2 recording sessions and 1 animal in the hippocampus. n = 12 episodes from 2 recording sessions and 1 animal in the hypothalamus. n = 12 episodes from 2 recording sessions and 1 animal in the pons) (Kruskal-Wallis: F(11, 228) = 109.5, *p* < 0.05 in the cortex; Kruskal-Wallis: F(11, 60) = 55.5, *p* < 0.05 in the hippocampus. Kruskal-Wallis: F(11, 132) = 46.8, *p* < 0.05 in the hypothalamus; Kruskal-Wallis: F(11, 132) = 61.2, *p* < 0.05 in the pons, followed by multiple comparisons with the Bonferroni test: *p* < 0.05 vs the fourth epoch immediately before state transition).

We also performed correlation analysis to investigate the association between episode duration of REM and changes in Ca^2+^ concentration among the brain regions (Fig. 3D). Changes in cortical and hippocampal Y/C ratios were not found to correlate with episode duration of REM sleep (n = 22 episodes from 6 recording sessions and 3 animals in the cortex. n = 6 episodes from 2 recording sessions and 1 animal in the hippocampus) (*r* = 0.28, *p* = 0.21, NS in the cortex; *r* = 0.61, *p* = 0.20, NS in the hippocampus). In the hypothalamus and pons, however, episode duration of REM sleep and changes in Y/C ratios showed a significant negative correlation (n = 11 episodes from 2 recording sessions and 1 animal in the hypothalamus. n = 5 episodes from 2 recording sessions and 1 animal in the pons) (*r* = −0.67, *p* < 0.05 in the hypothalamus; *r* = −0.97, *p* < 0.05 in the pons). These results suggest that the dynamics of astrocyte Ca^2+^ concentration varies depending on the brain region. A longer episode duration of REM sleep does not induce a further decrease in Ca^2+^ level in various regions of the brain, in this case in the cortex and hippocampus.

Next, we analyzed the correlation between electrophysiological features from cortical EEGs and EMGs among the sleep/wakefulness states, and astrocyte Ca^2+^ concentrations, as in Figure 2. As we compared cortical EEGs in this experiment, we focused on analyzing its correlation with the Ca^2+^ dynamics of cortical astrocytes (6 recording sessions and 3 animals). During wakefulness, Y/C ratios showed a weak negative correlation with alpha and beta waves and a weak positive correlation with gamma waves (Fig. 4A, 4D, and 4G). It was reported that an increase in the power value of delta and alpha waves and a decrease in the power value of gamma waves indicate a decrease in the arousal level during wakefulness (McGinley et al., 2015). The cortical astrocyte Ca^2+^ level appears to be low when the mice are in quiet wakefulness. During NREM sleep, the Y/C ratios were positively correlated with delta waves and EMG rms, and were negatively correlated with theta, alpha, and beta waves, consistent with those in the cerebellum (Fig. 4B, 4E, and 4H). During REM sleep, however, Y/C ratios demonstrated a positive correlation with delta and EMG rms, and a negative correlation with theta and alpha waves (Fig. 4C, 4F, and 4I). In this correlation analysis, minor but significant differences were observed also in the cerebellum. Compared with the results of the cerebellum, a tendency was observed of a correlation between astrocyte Ca^2+^ concentrations and electrophysiological features not only during sleep but also during wakefulness in the cortex.

### Region-specific Ca^2+^ dynamics in astrocytes

We further quantified the regional differences in Ca^2+^ signals, including in the cerebellum. First, we quantified Ca^2+^ dynamics during each episode by categorizing the episodes into 5 states, depending on the sleep/wakefulness state before and after each episode (Fig. 5A). Each episode was divided into 5 time-segments, and the average Ca^2+^ signal was calculated for each brain region (n = 6 recording sessions and 3 animals in the cortex. n = 2 recording sessions and 1 animal in the hippocampus. n = 4 recording sessions and 2 animals in the hypothalamus. n = 2 recording sessions and 1 animal in the pons. n = 5 recording sessions and 3 animals in the cerebellum). Based on the results of Figure 3 and Figure 5A, there are likely to be three clusters in quality. The first cluster is the cortex and hippocampus. Cortical astrocytes showed no significant Ca^2+^ concentration changes during the transition from wakefulness to NREM sleep, but showed a decrease in Ca^2+^ concentration at the time of NREM sleep transitioning to REM sleep, showing a trend similar to that of hippocampal astrocytes. In addition, no correlation was observed with duration of the REM sleep episode. The second cluster is the hypothalamus and pons. Astrocytes of both regions maintained their Ca^2+^ concentrations during NREM sleep, which decreased during REM. The third cluster is the cerebellum. Cerebellar astrocyte has characteristics of the other two clusters. Ca^2+^ changes associated with sleep/wakefulness states tend to resemble those of the cortex and hippocampus. However, as in the hypothalamus and brainstem, Ca^2+^ changes negatively correlated with duration of the REM episode.

**Figure 5.**
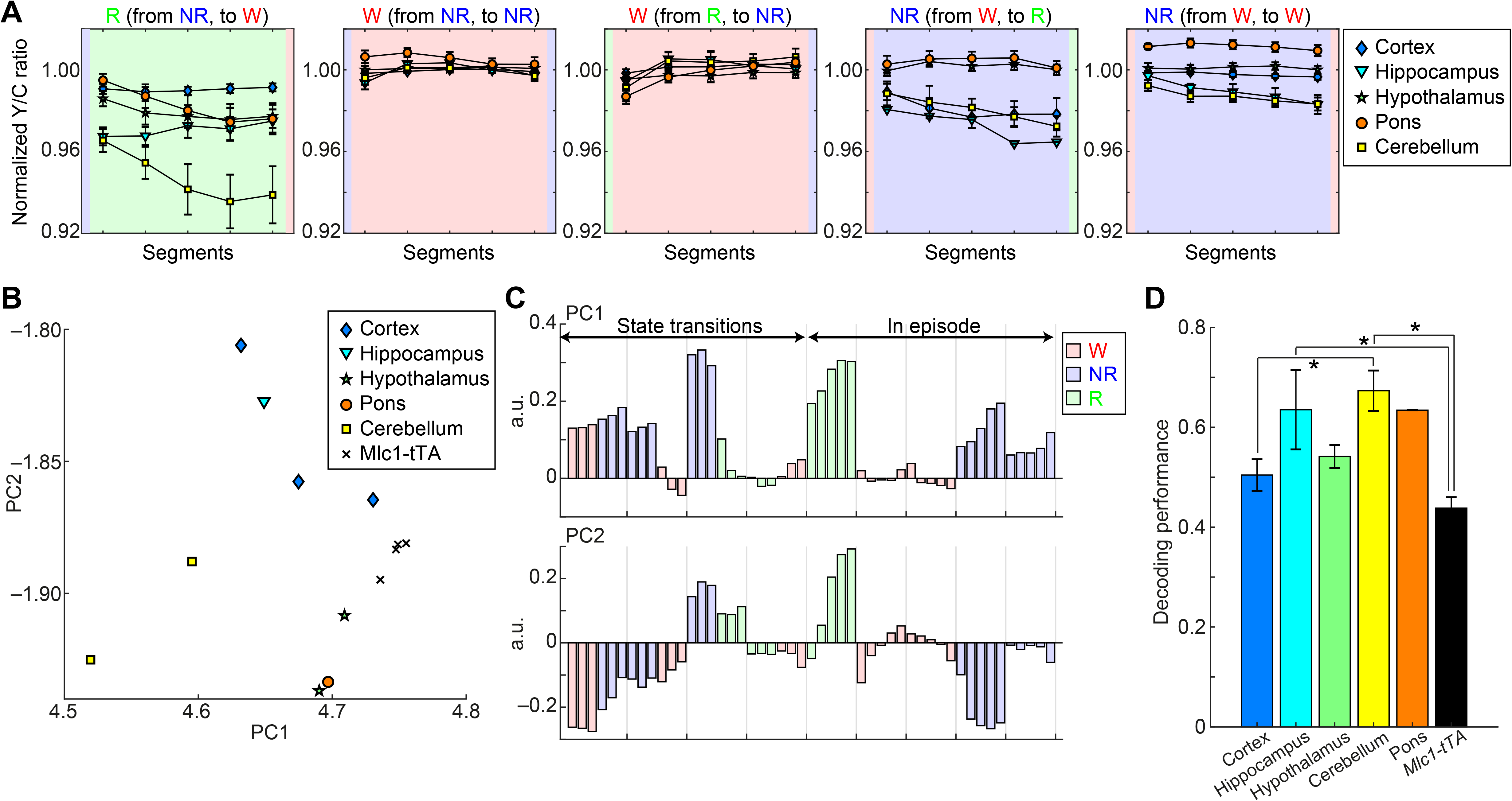
Dynamics of Ca^2+^ signals during different sleep/wakefulness states and different brain regions. (A) The mean profile of Ca^2+^ signals during time-normalized episodes. In each panel, the duration of each episode was segmented into 5 bins and the mean normalized Y/C ratios were computed in various brain regions. (B) PCA of the dynamics of Ca^2+^ signals in different brain regions in each mouse. The first two principal components accounted for 88% of the total variance. Each dot represents a mouse. (C) The first two principal components. (D) Decoding performance of Ca^2+^ signals for different sleep/wakefulness states among various brain regions. *, *p* < 0.05, *F*(5, 17) = 6.10, one-way ANOVA with the post-hoc HSD test.

Next, to visualize and quantify state-dependent and region-specific Ca^2+^ dynamics in low dimensional space, PCA was performed by constructing a vector for each mouse with a mean state transition and episode (Fig. 5B and 5C). Recordings from the cerebellum and hypothalamus of *Mlc1*-tTA mice was used as the control (n = 8 recording sessions and 3 animals). In Figure 5B, each dot represents a mouse. The first two principal components accounted for 88% of the total variance. The first principal components (PC1) represents the components that classify the cerebellum, and the others, while the second principal components (PC2) represents the components that classify the cortex and hippocampus, and the others, showing a tendency to form region-specific clusters. There appeared to be three clusters, the first one is the cortex and hippocampus, the second one is the cerebellum, and the third is the hypothalamus and pons. Figure 5C represents the profiles of PC1 and PC2. Since PC1 tends to show positive values during sleep, it is expected that mice with higher Ca^2+^ concentrations during sleep show higher value of the horizontal axis in Figure 5B.

Next, we analyzed the extent to which astrocyte Ca^2+^ signals can predict the ongoing sleep/wakefulness state, and whether there are any differences in decoding performance among the brain regions. For this purpose, average Ca^2+^ signals were computed in each epoch and a linear classifier was trained with 4-fold cross validation (Tsunematsu et al., 2020). The decoding performance of the hippocampus and cerebellum based on astrocyte Ca^2+^ signals were significantly higher than that from *Mlc1*-tTA mice (Fig. 5D; *p* < 0.05, one-way ANOVA). This result indicates that hippocampal and cerebellar astrocyte activity demonstrate sleep/wakefulness state-dependency. In addition, we also clarified the differences in decoding performance among different brain regions. Decoding performance of the cerebellum was significantly higher than that of the cortex (Fig. 5D; *p* < 0.05, one-way ANOVA). Thus, the physiological role of astrocytes in sleep/wakefulness might vary depending on the brain region.

## Discussion

In the present study, we analyzed astrocyte Ca^2+^ dynamics during different sleep/wakefulness states among various brain regions. Our study demonstrated that astrocyte Ca^2+^ concentration decreased during sleep, reached a minimum during REM sleep, and increased at the same time as wakefulness in all brain regions that we recorded. Further analyses indicated that there are at least three astrocyte clusters depending on the brain region. Although the association between changes in astrocyte Ca^2+^ concentrations and sleep/wakefulness were generally consistent among the brain regions unlike neural activity, we have found that Ca^2+^ dynamics varied depending on the brain region.

### Technical considerations compared with other similar studies

Several recent studies measuring astrocyte Ca^2+^ concentrations during the various sleep/wakefulness states have been reported (Bojarskaite et al., 2020; Ingiosi et al., 2020). These results are in very good agreement with our results showing that astrocytes Ca^2+^ concentrations/signals decrease during sleep and increase during wakefulness. However, there are several technical differences between the previous studies and our present study.

The first difference is the Ca^2+^ sensor that was used. We used the fluorescence resonance energy transfer (FRET)-based, ultrasensitive genetically encoded Ca^2+^ indicator YCnano50 to measure intracellular Ca^2+^ concentrations in astrocytes (Horikawa et al., 2010). On the other hand, other studies used GCaMP6f (Chen et al., 2013). YCnano50 has high Ca^2+^ affinity (Kd = 50 nM) compared with GCaMP6f (Kd = 375 nM). In addition, the ratiometric property of YCnano50 enables the reduction of motion artifacts during *in vivo* optical imaging. Because of the ability of YCnano50 to detect subtle basal changes in the Ca^2+^ concentrations of astrocytes, our study has clarified the distinct Ca^2+^ concentration differences between NREM and REM sleep.

The second difference is the method of expression of the Ca^2+^ sensor. Previous studies expressed the sensor using an adeno-associated virus, whereas we used *Mlc1*-tTA; TetO-YCnano50 bigenic mice to express the Ca^2+^ sensor in an astrocytes-specific manner. We previously demonstrated that *Mlc1*-tTA; TetO-YCnano50 mice together with the knockin-mediated enhanced gene expression by improved tetracycline-controlled gene induction (KENGE-tet) system enables extensive expression of YCnano50 in astrocytes in the brain (Tanaka et al., 2012; Kanemaru et al., 2014). In addition, the use of genetically modified mice rather than virus infection enabled us to record various regions of the brain, and the expression level and expression pattern of YCnano50 were constant among mice and hence consistent data could be obtained.

The third is the optical imaging approach that was used. As we recorded Ca^2+^ levels in astrocytes using the fiber photometry system, it was possible to analyze deeper regions of the brain, such as the hypothalamus and pons. However, as our method only enables the measurement of the sum of changes in Ca^2+^ concentration of cells at the tip of the inserted optical fiber, the behavior at the single-cell level remains unknown, as for measurements by two-photon microscopy. In addition, it was not possible to clarify whether or not subcellular regions of the astrocytes, such as the soma and distal processes, behave differently.

Taking advantage of our method, we succeeded in showing the existence of three astrocyte clusters and compared them among various brain regions.

### Possible mechanisms regulating astrocyte Ca^2+^ concentrations

By measuring changes in astrocyte Ca^2+^ levels during various sleep/wakefulness states, we observed similar Ca^2+^ concentration changes in different brain regions. This was an interesting result that was inconsistent with neural activity. For instance, cortical neurons are activated during wakefulness and REM sleep (Vyazovskiy et al., 2009; Watson et al., 2016; Niethard et al., 2017), hippocampal neurons fire less during REM sleep (Grosmark et al., 2012; Miyawaki and Diba, 2016), and an increase in the firing rate of neurons in the brainstem is observed during REM sleep (McCarley and Hobson, 1971; Hobson et al., 1975; Weber et al., 2015; Tsunematsu et al., 2020). Thus, it has been reported that neural activity patterns vary depending on the brain region.

It has been reported that Ca^2+^ concentrations are affected by G-protein coupled receptors (GPCRs) expressed in astrocytes via neurotransmitter release that accompanies neural activity. In general, the activation of astrocyte GPCRs increases astrocyte Ca^2+^ levels (Cornell-Bell et al., 1990; Takata et al., 2011; Jacob et al., 2014; Corkrum et al., 2020), although in some instances a decrease in Ca^2+^ level has been reported (Jennings et al., 2017). However, we observed global Ca^2+^ concentration changes, and therefore it is possible that a neurotransmitter that changes throughout the brain during different sleep/wakefulness states controls astrocyte Ca^2+^ concentration. Noradrenaline has been reported to increase astrocyte Ca^2+^ levels (Bekar et al., 2008; Paukert et al., 2014; Oe et al., 2020). Noradrenergic neurons located in the locus coeruleus (LC) project to the entire brain. Changes in firing rates of noradrenergic neurons showed a similar pattern to the Ca^2+^ dynamics of astrocytes (Takahashi et al., 2010; Tsujino et al., 2013). Taken together, noradrenaline released form noradrenergic neurons during wakefulness might increase astrocyte Ca^2+^ concentrations throughout the brain. In addition, considering that astrocyte Ca^2+^ levels gradually decrease during REM sleep, noradrenaline might also act as a volume-transmitter, because microdialysis studies have reported that the concentration of noradrenaline in the brain decreases during sleep (Park, 2002; Bellesi et al., 2016). It has been reported that not only noradrenaline but also glutamate, acetylcholine, and gamma-aminobutyric acid (GABA) increase the concentration of astrocyte Ca^2+^ levels (Cornell-Bell et al., 1990; Kang et al., 1998; Araque et al., 2002; Sun et al., 2013; Perea et al., 2016). Thus, the effects of these neurotransmitters should not be ignored and should be elucidated in the near future.

The activity of monoaminergic neurons, including noradrenergic neurons, alters depending on the sleep/wakefulness state. On the other hand, as neuropeptides, such as orexin and melanin-concentrating hormone (MCH), are known to be important for regulation of the sleep/wakefulness state (Chemelli et al., 1999; Sakurai, 2007; Tsunematsu et al., 2014), it is necessary to further analyze the association between neuropeptides and astrocyte Ca^2+^ dynamics.

### Possible heterogeneity of astrocytes among various brain regions

A surprising implication in this study is that astrocytes in different brain regions may have different functions in sleep/wakefulness. PCA classified astrocytes into the following three clusters: cluster 1, the cortex and hippocampus; cluster 2, the cerebellum; and cluster 3, the hypothalamus and pons. These clusters imply functional differences.

Astrocytes have recently been clarified to be a heterogeneous population. Gene expression analyses have demonstrated that astrocyte gene expression patterns differ among and within brain regions and can be classified into several types (Chai et al., 2017; Morel et al., 2017; Zeisel et al., 2018; Batiuk et al., 2020; Bayraktar et al., 2020; Lozzi et al., 2020). PCA of the gene expression patterns indicated that cortical and hippocampal astrocytes are transcriptionally close and form similar clusters (Morel et al., 2017; Lozzi et al., 2020). In contrast, Bergmann glial cells are a type of cerebellar astrocyte with unique morphological and transcriptional characteristics although there are other glial cells, velate astrocytes, in the cerebellum. Bergmann glial cells normally express Ca^2+^-permeable AMPA receptors composed of the GluA1 and GluA4 subunits (Saab et al., 2012), implying that they have different intracellular Ca^2+^ dynamics compared with other glial cells. These previous reports were consistent with our results based on PCA. Our results may explain part of the differences in the functions of astrocytes depending on the brain region, as well as the differences in their cellular transcriptomes, although it is still early to make a conclusion as the number of mice analyzed in our study was limited. In addition, heterogeneous gene expression patterns have been found within brain regions, for example, in the cortical layer (Bayraktar et al., 2020). Furthermore, detailed studies will be necessary to analyze the Ca^2+^ dynamics in astrocytes within the same brain regions during different sleep/wakefulness states.

The decoding performance for ongoing sleep/wakefulness states was also significantly different depending on the brain region. Decoding performance was significantly higher in the hippocampus and cerebellum of *Mlc1*-tTA; TetO-YCnano50 bigenic mice than control *Mlc1*-tTA monogenic mice. In other words, our results suggest that astrocyte Ca^2+^ dynamics in the hippocampus and cerebellum might not only show state-dependent fluctuation, but may also contribute to the control of the sleep/wakefulness state itself. In the future, it will be necessary to clarify the functions of astrocytes in different sleep/wakefulness states. Further research should also be carried out with the caveat that astrocyte functions might vary among different brain regions.

## Acknowledgements

This work was supported by PRESTO from Japan Science and Technology Agency (JST) (grant no.: JPMJPR1887) and by JSPS KAKENHI Grant Number 20H05047 to T.T. and Grant-in-Aid for Scientific Research (B) (19H03338), Grant-in-Aid for Challenging Exploratory Research (18K19368), Toray Science Foundation to K.M. We thank A. Utsumi for assistance with the data analyses, and Dr. Helena Akiko Popiel for English language editing of the manuscript.

